# Phenotype-guided subpopulation identification from single-cell sequencing data

**DOI:** 10.1101/2020.06.05.137240

**Authors:** Duanchen Sun, Xiangnan Guan, Amy E. Moran, David Z. Qian, Pepper Schedin, Andrew Adey, Paul T. Spellman, Zheng Xia

## Abstract

Single-cell sequencing yields novel discoveries by distinguishing cell types, states and lineages within the context of heterogeneous tissues. However, interpreting complex single-cell data from highly heterogeneous cell populations remains challenging. Currently, most existing single-cell data analyses focus on cell type clusters defined by unsupervised clustering methods, which cannot directly link cell clusters with specific biological and clinical phenotypes. Here we present Scissor, a novel approach that utilizes disease phenotypes to identify cell subpopulations from single-cell data that most highly correlate with a given phenotype. This “phenotype-to-cell within a single step” strategy enables the utilization of a large amount of clinical information that has been collected for bulk assays to identify the most highly phenotype-associated cell subpopulations. When applied to a lung cancer single-cell RNA-seq (scRNA-seq) dataset, Scissor identified a subset of cells exhibiting high hypoxia activities, which predicted worse survival outcomes in lung cancer patients. Furthermore, in a melanoma scRNA-seq dataset, Scissor discerned a T cell subpopulation with low *PDCD1*/*CTLA4* and high *TCF7* expressions, which is associated with a favorable immunotherapy response. Thus, Scissor provides a novel framework to identify the biologically and clinically relevant cell subpopulations from single-cell assays by leveraging the wealth of phenotypes and bulk-omics datasets.

## Introduction

Single-cell sequencing technologies are revolutionizing biomedical research and clinical practice by enabling the comprehensive characterization of cells from complex tissues^1-3^. In contrast to bulk data that measures the averaged properties of whole tissue, single-cell sequencing allows the identification of cell types, states and lineages of different cell subpopulations in a heterogeneous tissue ecosystem^4-6^. To recognize critical subpopulations from single-cell data, the standard approach is to perform unsupervised clustering to define cell clusters, inspect marker genes of each cluster, and assess the enrichment of the marker genes in known cell types and pathways^7-9^ to evaluate the importance of each cell cluster. However, identifying cell subpopulations that drive phenotypes, such as disease stage, tumor subtypes, treatment response and survival outcome, is of the greatest interest, since it will facilitate cell-type targeted therapies as well as prognostic biomarker discoveries in diseases^4, 10^. Unfortunately, single-cell technology is not practical in large cohort and most single-cell experiments involve less than twenty patient samples^3, 11, 12^, which lack the statistical power to identify the cell subpopulations driving the phenotype of interest.

Meanwhile, valuable clinical phenotype information is widely available from big data consortia like The Cancer Genome Atlas (TCGA) through a decade long collection of clinicopathologic annotations^13, 14^. Clinical phenotype information is primarily collected on bulk tissue samples, especially in the form of formalin-fixed-paraffin-embedded (FFPE) samples, which are not feasible for single-cell profiling. Therefore, there is an unmet need to leverage such widely accessible and valuable phenotype information to guide cell subpopulation identification from single-cell data.

To the best of our knowledge, there is no bioinformatics tool that uses phenotypes to guide the identification of key cell subpopulations in a unified framework. Therefore, in this study, we introduce **S**ingle-**C**ell **I**dentification of **S**ubpopulations with bulk RNA-**S**eq phenotype c**OR**relation (Scissor) (Fig.1). By leveraging bulk data and phenotype information (Fig. 1a), this algorithm automatically selects a subset of cells from the single-cell data that is most responsible for the differences of phenotypes (Fig. 1b). Since a subpopulation with a small number of cells among the thousands of cells may drive the phenotype of interest^15^, we employ a graph-regularized sparse regression model to select the cell subsets that are important for the given phenotype with high confidence. According to the signs of the estimated regression coefficients, the selected cells are indicated as Scissor positive (Scissor+) cells and Scissor negative (Scissor-) cells (Fig. 1c), which are positively and negatively associated with the given phenotype, respectively. The non-selected cells with coefficients 0 are indicated as background cells (Fig. 1c). The selected cells will be further analyzed in terms of signature genes and functional enrichment analysis (Fig. 1d).

**Figure 1.**
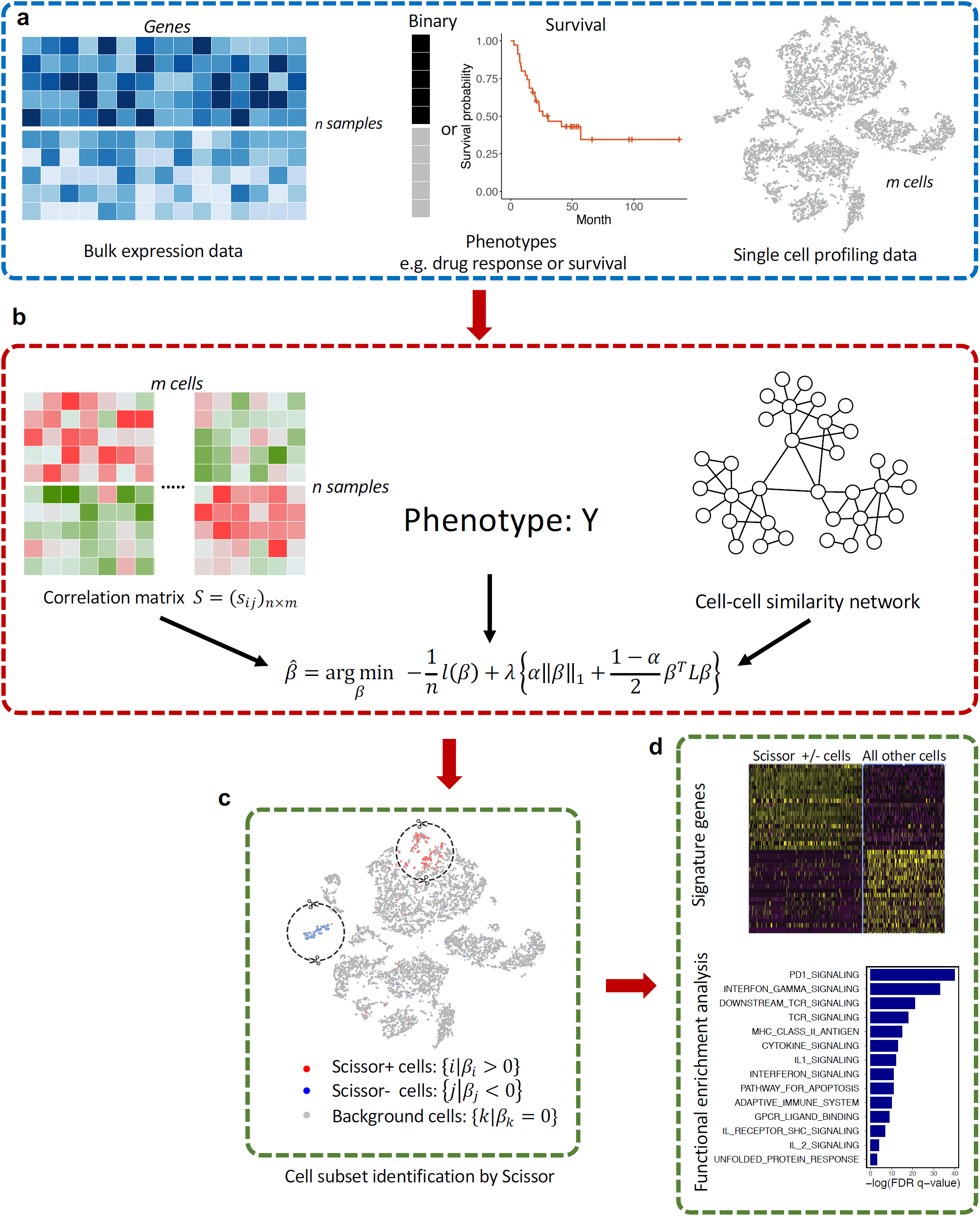
The workflow of phenotype guided cell subpopulation identification method Scissor. (**a**) The inputs for Scissor: a bulk expression matrix, a corresponding phenotype such as drug response or clinical information, and a scRNA-seq data needs to be analyzed. (**b**) Scissor calculates a correlation matrix and a cell-cell similarity network based on input sources, which are further integrated with the phenotype into a network regularized sparse regression model to select the most relevant cell subpopulations. (**c**) The selected cells by Scissor with red and blue dots indicating the cells positively or negatively associated with the phenotype. (**d**) The selected cells by Scissor can be further investigated by downstream analyses.

The novelty of Scissor is that it utilizes phenotype information from bulk RNA-seq data to identify the most highly disease-relevant cell subsets. When we applied Scissor to real datasets, the selected cells exhibited high specificity of association with the phenotype of interest. Our studies suggest that Scissor is a promising tool to explore and interpret single-cell data from a new perspective, which can shed fresh light on disease mechanisms and improve the diagnosis and treatment of diseases.

## Results

### Scissor identified cell subpopulations related to the tumor and normal phenotypes from scRNA-seq

We first applied Scissor to a lung cancer scRNA-seq data that included tumor cells and cells from the tumor microenvironment with cell types defined in the original paper^11^ (Fig. 2a). In order to demonstrate the effectiveness of our algorithm, we utilized the phenotype defined as the tumor and normal from TCGA lung adenocarcinoma (LUAD) 577 bulk RNA-seq samples^16^ to guide our Scissor analysis. We expected that by using these data, we would be able to infer cells that were most highly associated with the cancer or normal phenotype in this heterogeneous single-cell dataset. Because of the binary phenotype settings of this application, where samples with a phenotype indicator value 1 correspond to tumor bulk samples, the Scissor+ cells should be associated with cancer cells and the Scissor-cells should be associated with the normal phenotype. Among 29,888 cells from different cell types (Fig. 2a), 361 Scissor+ cells and 534 Scissor-cells were selected by Scissor with the default parameters, which were related to the tumor and normal phenotypes with high confidence (Fig. 2b). As anticipated, over 98% of Scissor+ cells were verified to be malignant cells and the remaining were T cells and B cells (Fig. 2c) based on the cell type annotated in the Ambrechts *et al*. study^11^. Such a high proportion of cancer cells in Scissor+ cells cannot be selected by chance (Fig. 2d, hypergeometric test *p*<2e-16), which confirmed that Scissor can identify the cells associated with the phenotypes of interest. As for Scissor-cells, the cell types were relatively more balanced than Scissor+ cells since it was designed to correlate with more diverse non-malignant cell types (Fig. 2c). Myeloid cells and alveolar cells were two main selected cell types, which account for 42.3% and 36.9% of total Scissor-cells, respectively. All cell types in Scissor-cells, especially the alveolar cells, are important cell types of normal lungs^17^. Thus, we demonstrated that Scissor can precisely identify the most phenotype-associated cells from single-cell data with the guidance of phenotype information from bulk data.

**Figure 2.**
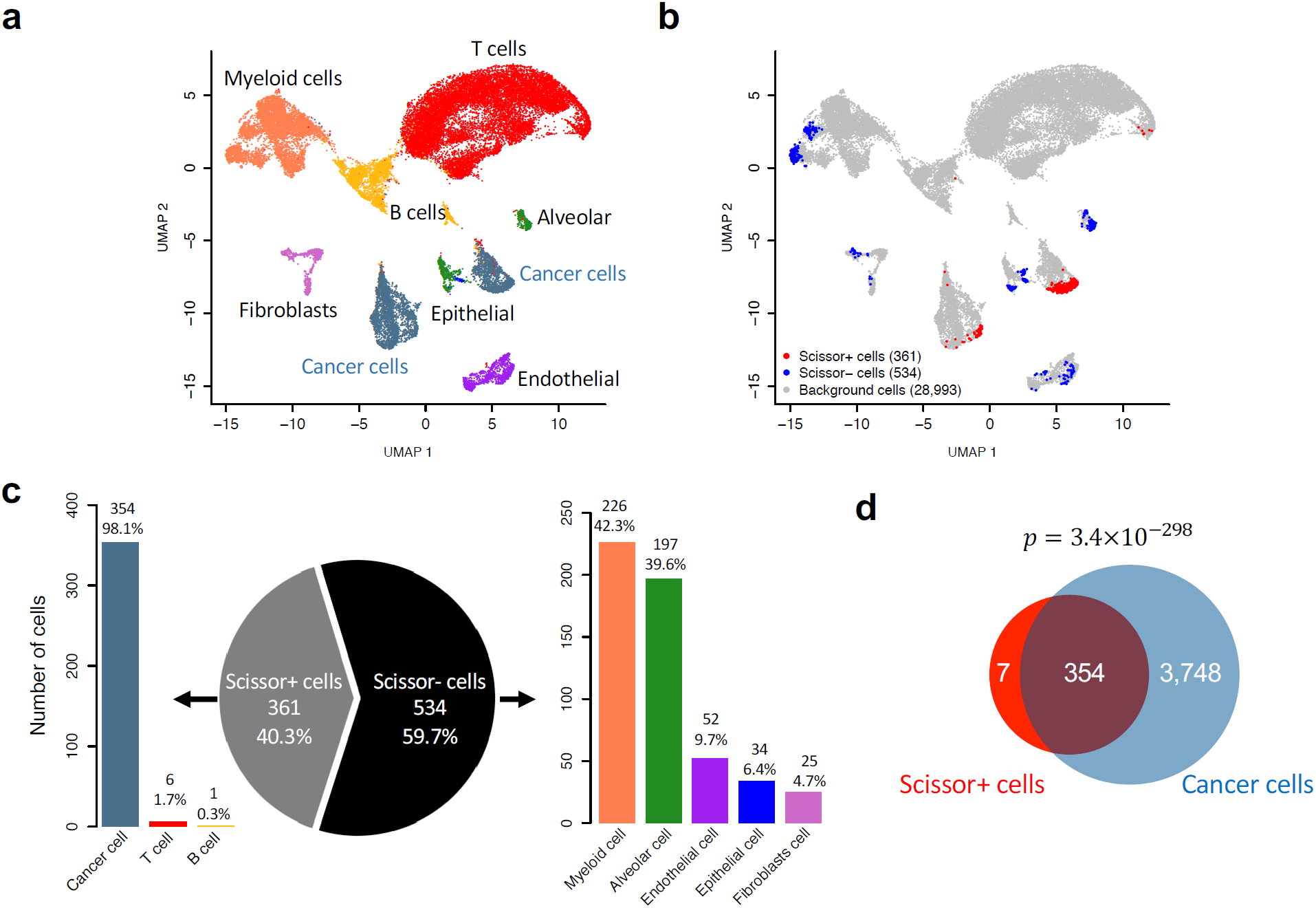
Scissor identified subpopulations related to the tumor and normal phenotypes from scRNA-seq. (**a**) UMAP visualization for 29,888 LUAD cells^11^. The color encoding the cell types defined in the original paper^11^. (**b**) Scissor selected cells as indicated by the red and blue dots on the same UMAP plot. (**c**) The pie chart of Scissor selected cells with corresponding bar-plots showing the detailed constitution in each portion. (**d**) The Venn diagram showed a significant overlap between Scissor+ cells and cancer cells. The statistical *p* value was calculated by hypergeometric test.

### Scissor identified a hypoxic subpopulation of lung cancer cells correlated with worse survival

Cancer cells are heterogeneous and include subpopulations such as cancer stem cells, which are known to drive tumor progression, drug resistance and poor outcomes^18-20^. Therefore, we applied Scissor, guided by the TCGA-LUAD 471 bulk RNA-seq samples and their corresponding survival information^16^, to identify aggressive cancer cell subpopulations within 4,102 cancer cells from a lung cancer scRNA-seq dataset^11^. These cells were separated into 12 clusters (Fig. 3a), which demonstrated the heterogeneous nature of the cancer cells. Out of 205 Scissor selected cells, 201 Scissor+ cells were associated with **W**orse **S**urvival (defined as Scissor_WS cells thereafter) and only 4 Scissor-cells were associated with good survival (Fig. 3b). The Scissor_WS cells were mainly from clusters 1 and 3 (Fig. 3c). To understand the underlying transcriptional patterns of Scissor_WS cells, we compared the gene expressions of those cells with all other cells (consisting of Scissor- and background cells) following the criteria outlined in our methods section. As a result, 23 up-regulated genes and 205 down-regulated genes were differentially expressed in Scissor_WS cells over all other cells, respectively (Fig. 3d, Supplementary Table 1,2). Interestingly, we found that multiple important hypoxia-related genes (*GAPDH, ENO1, TPI1, LDHA, ALDOA, PGK1* and *FAM162A*, Fig. 2e) were among the above 23 overexpressed genes. Functional enrichment analysis also confirmed that hypoxia-related pathways, such as glycolysis and glucose metabolism processes, were activated in Scissor_WS cells (Fig. 3f). Consistently, motif analysis revealed that *HIF1A* binding motif was the most enriched one in the 23 up-regulated genes (Supplementary Table 3), which is a key mediator of the cellular response to lowered oxygen levels and plays critical roles to adapt the metabolism of cancer cells to a hypoxic microenvironment^21^. Therefore, the hypoxic Scissor_WS cell population might drive the worse clinical outcomes since oxygen-starved cancer cells have survival advantage to elevate tumor progression^22^.

**Figure 3.**
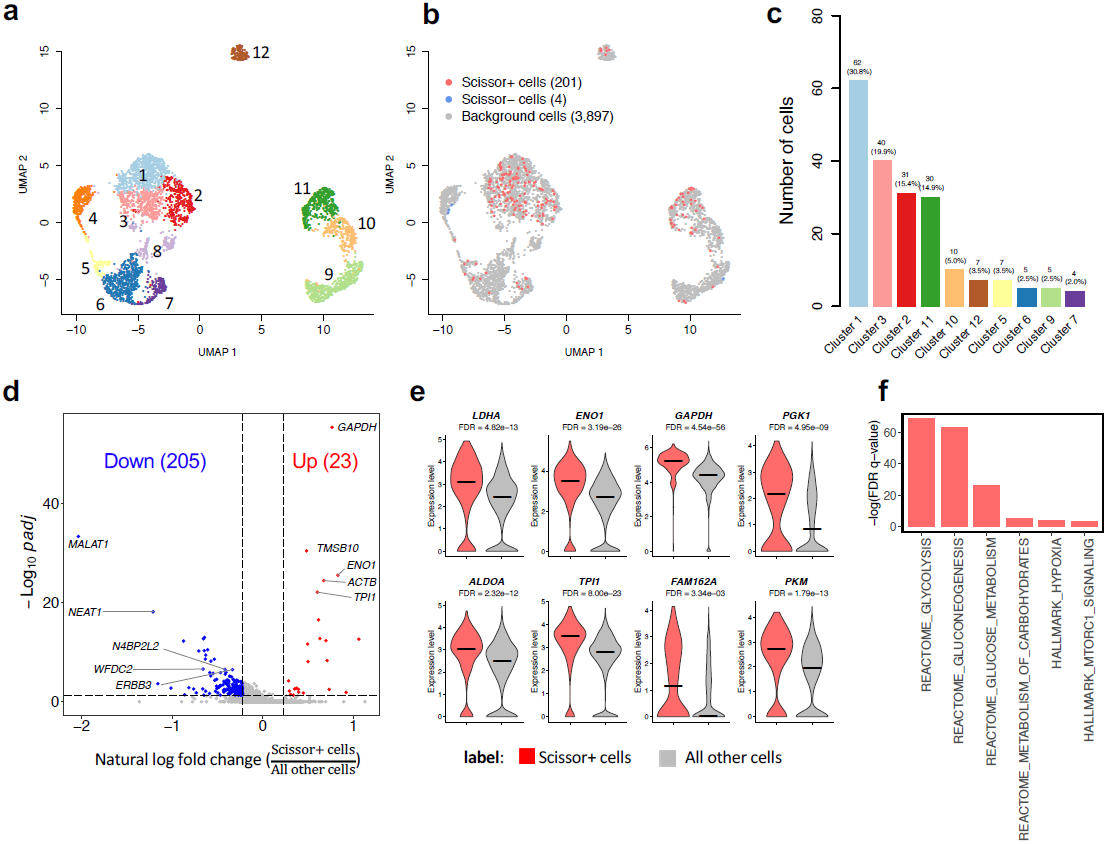
Scissor identified a hypoxic subpopulation of lung cancer cells guided by the TCGA-LUAD survival outcomes and bulk expressions. (**a**) UMAP visualization for 4,102 LUAD cancer cells. The cell clusters were inferred by Seurat^31^. (**b**) UMAP visualization for Scissor selected cells. The red and blue dots are Scissor+ (worse survival) and Scissor-(good survival) cells, respectively. (**c**) The bar-plot shows the detailed constitution in Scissor+ cells in different clusters. (**d**) Volcano plot representation of differential expression analysis of genes in the Scissor+ cells versus all other cells (including Scissor-cells and background cells). Red and blue points mark the genes with significantly increased or decreased expression respectively in Scissor+ cells compared to other cells (FDR <0.05 and natural log fold change > 1.25). The two vertical dashed lines represent ±ln(1.25) fold-changes in gene expression and the horizontal dashed line denotes FDR cutoff 0.05. (**e**) The violin plots of expression levels of selected up-regulated genes in Scissor+ cells. The FDR was calculated by Wilcoxon Rank Sum test in Seurat. (**f**) Enrichment bar plot of selected hypoxia-related pathways in Reactome and Hallmark domains.

### The subpopulation identified by Scissor from lung cancer cells revealed a clinically relevant signature

In the previous application, Scissor identified an aggressive cell subpopulation (Scissor_WS) that was associated with the worse survival and can be characterized by the overexpression of hypoxia-related genes. To further examine the clinical relevance of the above 23 overexpressed genes (defined as lung cancer signature), we chose six independent gene expression datasets of lung cancer collected in a public website^23^ (see Methods for more details). We used Gene Set Variation Analysis (GSVA)^24^ to calculate the signature score of the lung cancer signature in each sample, which can be used to stratify samples for survival analysis. Indeed, we found that in 5 out of 6 datasets, the patients with high signature scores had significantly worse survival time than the patients with low signature scores (Fig. 4a, Supplementary Figure 1, Supplementary Table 4), which indicated that the lung cancer signature derived from the cell subpopulation identified by Scissor was associated with worse survival and could serve as a biomarker for poor clinical prognosis.

**Figure 4.**
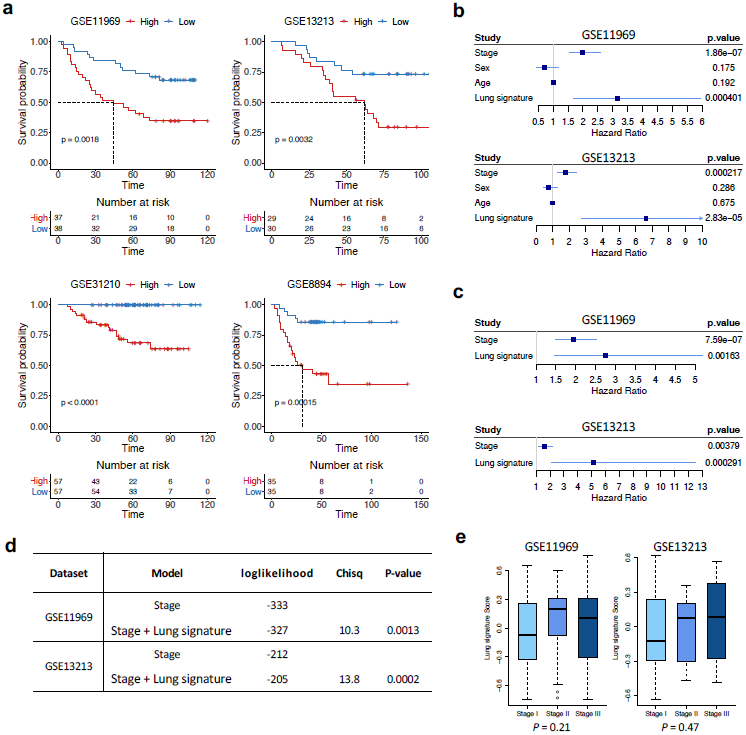
Clinical utility of the lung cancer signature derived from the subpopulation identified by Scissor. (**a**) Lung cancer signature relates to prognostic information on 4 independent lung cancer datasets. For each dataset, samples were divided into two groups based on the quantile values of lung cancer signature and Kaplan-Meier survival curve was drawn for each group. A corresponding table is also attached to display the number of alive samples at given time points and the statistical *p* value was calculated by log-rank sum test. (**b**) Forest plots showed the hazard ratio and 95% confidence interval for lung cancer signature and three additional clinical features according to univariate Cox model. (**c**) Forest plots show the hazard ratio and 95% confidence interval for lung cancer signature and stage information according to multivariable Cox model. (**d**) The ANOVA deviance table fitted by cox models with or without lung cancer signature. The likelihood ratio test was used to test for a significant difference in model fit between the model with tumor stage alone and the model including both tumor stage and lung signature. (**e**) Distributions of the lung cancer signature scores in terms of tumor stages. The statistical *p* values were calculated by Kruskal-Wallis test.

Among the six chosen datasets, two of them have additional clinical features like tumor stage. We thereby investigated whether our lung cancer signature can add additional prognostic power beyond clinical features. To achieve this, we examined pathological stage, sex and age at diagnosis and our lung cancer signature in these two datasets. First, a univariate Cox proportional hazard model was adopted to examine the association between survival time and these features. We found that only pathological stage and our signature were significantly associated with survival (Fig. 4b). Second, a multivariable Cox model was used to further evaluate the contributions of these two measurements to the survival analysis. We found that our signature remained statistically significant in both datasets after adjusting for tumor stage (Fig. 4c). Interestingly, combining our signature with tumor stage significantly improved the survival prediction power compared to using tumor stage alone (Fig. 4d, likelihood ratio test *p*=0.0013 and 0.0002, respectively). We further confirmed that our signature did not have statistical differences between different stages (Fig. 4e, Kruskal-Wallis test *p*=0.21 and 0.47, respectively), which indicated that our signature was independent of tumor stage. These data indicated that our lung cancer signature derived from Scissor analysis was complementary to tumor stage and could provide additional clinical power to improve prognostic evaluation of LUAD. In summary, these findings suggested that Scissor_WS cells with high hypoxia activity might drive LUAD progression and thereby conferred poor outcomes to patients whose tumors contained significant numbers of such cells. Thus, our Scissor analysis identified a cell subpopulation (Scissor_WS) from the LUAD scRNA-seq data that was associated with worse survival outcomes, and the lung cancer signature derived from Scissor_WS cells might help improve clinical prognosis as well as provide potential drug targets for further investigation^25^.

### Scissor identified a PD1 low subpopulation of T cells associated with response to melanoma immunotherapy

Single-cell sequencing is becoming a key player in immunology research especially the cancer immunotherapy field, by profiling different immune cell populations^2^. Immune checkpoint therapy by blocking CTLA4, PD-1 and PD-L1 has achieved exciting results in a wide variety of cancers^26, 27^. However, only a small portion of patients can benefit from current checkpoint inhibition (CPI). Thus, understanding the mechanism underlying CPI response is critical to improving the efficacy of these therapies in a broader patient population. Here, by integrating immunotherapy response information from public bulk RNA-seq studies, we performed Scissor analysis on a melanoma scRNA-seq dataset^28^ to identify T cell subpopulation that is related to CPI response even this scRNA-seq dataset itself lacking immunotherapy information.

Based on the established role of T cells in immunotherapy response, we used Scissor to analyze 1,894 T cells from the metastatic melanoma tumor microenvironment^28^ (Supplementary Figure 2a). To assist Scissor analysis, we collected the RNA-seq expression profiles of 70 melanoma patients with known immunotherapy response information from two studies^29, 30^ (Supplementary Figure 2b). In the standard scRNA-seq data analysis using Seurat package^31^, the 1,894 T cells were clustered into six cell populations (Fig. 5a). By performing Scissor on these cells, 105 out of 1,894 T cells were identified as Scissor+ cells, which were correlated with a **F**avorable immunotherapy **R**esponse and were defined as Scissor_FR cells thereafter (Fig. 5b). Our Scissor framework did not report any Scissor-cells that correlated with unfavorable immunotherapy responses. The 105 Scissor_FR cells were mainly residing in clusters 2 and 3 (Fig. 5c), indicating clusters 2 and 3 were more associated with effective responses than other clusters. To characterize the transcriptional identities of the Scissor_FR cells, we compared the gene expression of these cells with all other cells. In total, 17 up-regulated and 120 down-regulated differential expression genes were identified in Scissor_FR cells (Fig. 5d, Supplementary Table 5). The Scissor_FR cells that were associated with an effective CPI response had increased expression of genes linked to T cell memory (*CCR7* and *SELL*) and survival (*IL7R*)^32, 33^ as well as lower expression of inhibitory genes (*HAVCR2, LAG3, PDCD1, CTLA4*) and MHC II class genes (*HLA-DRB5, HLA-DRB1, HLA-DPA1, HLA-DQB2*, and *HLA-DRB6*) (Fig. 5d,e and Supplementary Table 5). The Scissor_FR cells also exhibited enhanced expression of transcript factor *TCF7* (encoding TCF1) that have been shown to associate with favorable outcomes in CPI treatment^34-36^ (Fig. 5e). The first most down-regulated gene in Scissor_FR cell population was *TOX*, which is involved in the CD8 T cell dysfunction^37, 38^ (Fig. 5e, Supplementary Table 5). Pathway enrichment analysis showed the Scissor_FR cells had higher TNFA signaling and lower activity of *CTLA4, PD1* signaling and LCMV/tumor exhaustion pathways (Fig. 5f).

**Figure 5.**
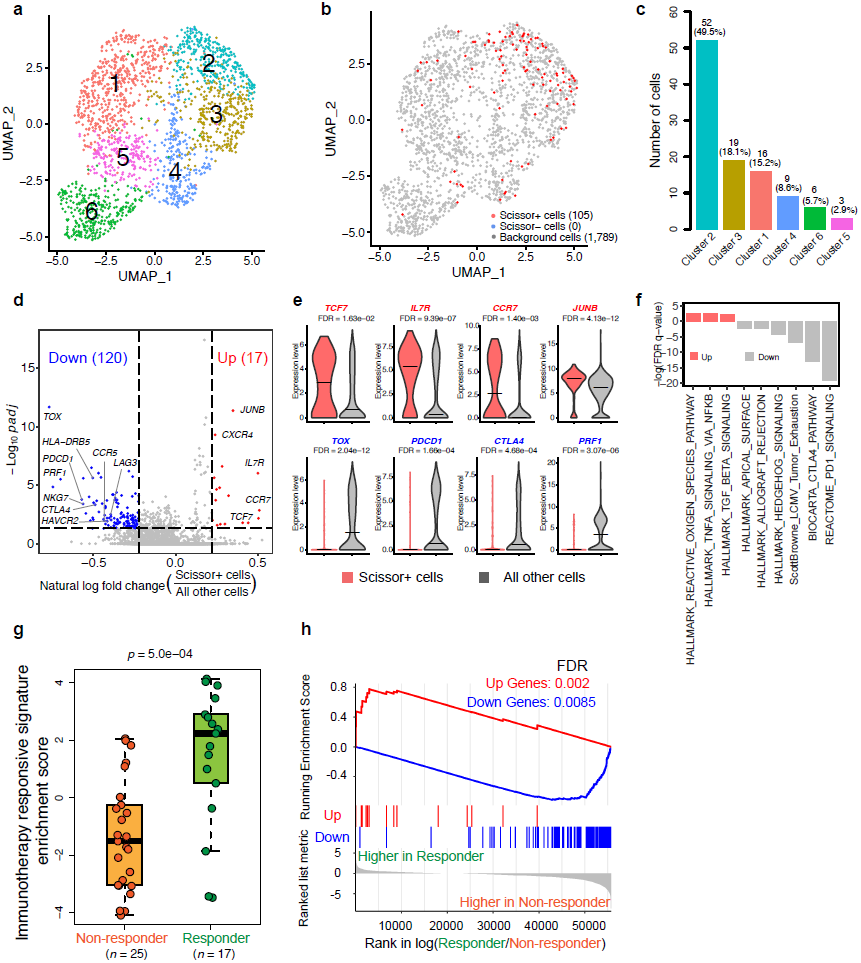
T cell subpopulation associated with effective immunotherapy response in melanoma. (**a**) UMAP plot showing six major T cell populations profiled from melanomas by scRNA-seq^28^. (**b**) The Scissor analysis selected cells correlated with immunotherapy responders or non-responders, which are highlighted in red or blue in the same UMAP plot, respectively. All other unselected cells are in gray. Scissor+ cells highlighted in red (Scissor_FR) are correlated with immunotherapy responders in melanomas. (**c**) Bar plot showing the distribution of selected Scissor+ cells across the six T cell populations. (**d**) Volcano plot of differential gene expressions of the Scissor+ cells compared with all other cells (including Scissor-cells and background cells). Genes highlighted in red and blue are the genes significantly up-regulated or down-regulated in the Scissor+ cells compared with all other cells, respectively (FDR <0.05 and natural log fold change > 1.25). (**e**) Violin plot showing the important immune signature genes of Scissor+ cells which are correlated with the immunotherapy responders. The FDR was calculated by Wilcoxon Rank Sum test in Seurat. (**f**) Bar plot showing the significantly enriched pathways in Scissor+ cells compared with all other cells (FDR <0.05). The up and down pathways were calculated by pathway analysis tool Camera^49^ (see methods). (**g**) Boxplot showing the enrichment score of immunotherapy responsive signature in the non-responders and responders from Sade-Feldman cohort. The statistical *p* value was calculated by Student’s *t*-test. (**h**) GSEA enrichment plot of the up and down genes in the responder vs non-responder comparison from the Sade-Feldman cohort.

Our Scissor analysis identified a subset of T cells that correlated with an effective immunotherapy response, suggesting that the 137 differential expression genes (defined as immunotherapy responsive signature) could be informative in the prediction of treatment success. To test this hypothesis, we applied this signature to an independent CPI melanoma dataset^34^. Indeed, the enrichment score of our signature was significantly higher in CPI responders than in non-responders (Fig. 5g, Student’s *t*-test *p*=5.0e-4). Additionally, the up-regulated and down-regulated genes in our immunotherapy responsive signature were significantly enriched in responders and non-responders of this independent dataset, respectively (Fig. 5h, Kolmogorov-Smirnov test FDR=0.002 and 0.0085, respectively). Our Scissor analysis thus identified a T cell subpopulation (Scissor_FR) that was associated with favorable immunotherapy response and the immunotherapy responsive signature derived from Scissor_FR cells could distinguish the responders from the non-responders to immunotherapy.

### Immunotherapy responsive signature genes reflect T cell States

Even though checkpoint blockade targets PD1 or CTLA4 expressed on exhausted T cells, our data showed that a cell subpopulation (Scissor_FR) with low *PDCD1*/*CTLA4* and high *TCF7* expression was associated with immunotherapy response. Since there is a strong relationship between immunotherapy response and T cell states^34^, we set out to explore the T cell states of our Scissor_FR subpopulation. To this end, we evaluated the enrichment scores of our immunotherapy responsive signature in five types of tumor-infiltrating lymphocytes (TILs) with distinctive differentiation states from a public dataset^38^. We found that the LAG3 low and PD1 low effector CD8 T cells (PD1_Lo_Eff) had the greatest enrichment scores of our signature, followed by the Naïve CD8, bystander TIL, LAG3 high/PD1 high effector CD8 (PD1_Hi_Eff) and exhausted CD8 T cells (Fig. 6a). Notably, our signature can significantly distinguish PD1_Lo_Eff, PD1_Hi_Eff and exhausted CD8 T cells from each other (Fig. 6a, Supplementary Figure 3). This observation indicated that our signature could reflect the state of the effector PD1_Lo_Eff TILs as well as distinguish different T cell differentiation states, which might help understand the T cell states of different TILs. Therefore, we further used our signature to perform a pseudotime analysis on the original melanoma TILs scRNA-seq dataset. The pseudotime trajectories revealed the relative orders of the 6 clusters defined in Fig. 5a, starting from cluster 2->3->4->5->6->1 (Fig. 6b). This result was consistent with the distribution of Scissor selected cells in different clusters, with the majority of Scissor_FR cells coming from the clusters 2 and 3 with low *PDCD1* expressions (Fig. 5c). Moreover, the PD1 low TILs also exhibit heterogeneity, containing memory-precursor-like and effector-like subsets, as reported by a recent study^39^. By comparing the enrichment score of our signature in the memory precursor and effector CD8 T cells from a public dataset^40^, we found that our signature was more enriched in the memory precursor CD8 T cells than the short-lived effector T cells (Fig. 6c, Student’s *t*-test *p*=0.02), which indicated that our selected responsive immunotherapy CD8 T subpopulation cells were more like PD1-low memory-precursor cells. The PD1-low memory-precursor-like T cells expressing *TCF7* have been shown to be associated with good immunotherapy response^39^. Collectively, our Scissor analysis of a scRNA-seq melanoma T cell dataset independently revealed a *PDCD1*/*CTLA4* low and *TCF7* high subpopulation of TILs whose distinct transcriptome was essential to the favorable response to immunotherapy. These data demonstrated that Scissor analysis of single-cell data is capable of identifying subpopulations correlated with specific phenotype even though the single-cell data itself has no such phenotype information.

**Figure 6.**
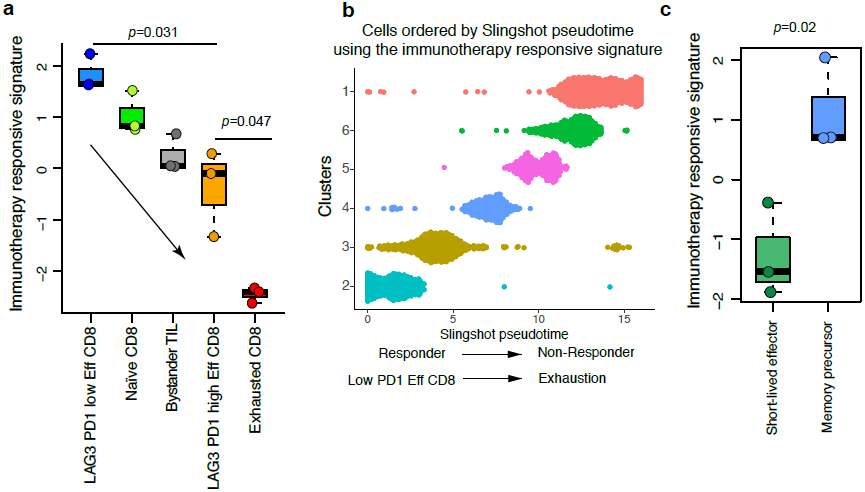
Immunotherapy responsive signature genes reflect T cell states. (**a**) Boxplot showing the enrichment score of immunotherapy responsive signature in 5 types of CD8 T cells sorted from a mouse liver tumor model^38^. Bystander TIL: Non-tumor-specific TCR_OT1_ cells isolated Day 20-21 from malignant liver lesions; Naïve CD8: naïve tumor-specific TCR SV40-I TILs; LAG3 PD1 low Eff CD8: PD1low/LAG3low/CD39low tumor-specific CD8 T cells isolated Day 8-9 from malignant liver lesions. LAG3 PD1 high Eff CD8: PD1high/LAG3high/CD39high TCR SV40-I CD8 T cells isolated Day 8-9 from malignant liver lesions; Exhausted CD8: TCR SV40-I CD8 T cells isolated Day 20-21 from malignant liver lesions. Student’s *t*-test was used to test the difference between the different groups. (**b**) Pseudotime analysis of the scRNA-seq of T cells from melanoma based on immunotherapy responsive signature. The clusters on the y-axis were defined as the same as Fig. 5a. (**c**) Boxplot showing the enrichment score of immunotherapy responsive signature in the memory precursor CD8 T cells and short-lived effector CD8 T cells^40^. The statistical *p* value was calculated by Student’s *t*-test.

## Discussion

Identifying the specific cell subpopulations that drive phenotypes, such as the poor prognosis and drug resistance, from single-cell assays is needed to advance the discovery of cellular targets for the treatment of disease. To this end, we present Scissor as a new computational tool for single-cell data analysis. In a unified mathematical model, Scissor utilizes the phenotype information from bulk data to select the cells that most highly correlate with a given phenotype from single-cell sequencing data.

The advantages of Scissor can be briefly summarized as follows: First, Scissor is a novel and effective computational method to identify cell subpopulations that are most highly associated with bulk phenotypes. The cells identified by Scissor exhibit different molecular properties from other cells, which involve crucial marker genes and biological processes of the given phenotype. Second, Scissor does not require any unsupervised clustering on single-cell data, which avoids subjective decisions of cell cluster numbers or clustering resolution^41^. Finally, Scissor provides a flexible framework for integrative analyses of single-cell data, bulk data and phenotype information. When applying Scissor to scRNA-seq datasets of lung cancer and melanoma, we identified a hypoxic subpopulation associated with worse survival in the TCGA-LUAD dataset, and a *PDCD1*/*CTLA4* low subpopulation of T cells in melanoma related to a favorable immunotherapy response. Thus, we demonstrated that Scissor can identify cell subpopulations associated with the phenotype information such as patient survival and response to therapy.

Scissor relies on a correlation matrix to quantify the dependence or similarity between single-cell data and bulk data. Other similarity measurement methods^42^ like entropy-based mutual information and hypothesis testing-based methods can also be used in Scissor, and may improve performance in some cases. Besides, although we demonstrated the integration of bulk RNA-seq with scRNA-seq data, Scissor can also be applied to other single-cell measurements like chromatin accessibility and DNA methylation^43^. For example, Scissor may help to identify the cell subpopulations based on chromatin accessibility by integrating single-cell ATAC-seq^22, 44^ of cancer cells and immune cells with the bulk ATAC-seq from TCGA^45^.

Overall, Scissor demonstrates promise for integrating single-cell data with phenotype information to dissect the clinically significant subsets from the heterogeneous cell populations. This strategy will boost biological discoveries and interpretation from a novel perspective in single-cell data analysis. We anticipate that Scissor will enable a broad application of widely available phenotype information on single-cell data analysis and help unravel the most disease-relevant subpopulations for cell-targeted therapies.

## Materials and Methods (Experimental Protocol)

### Single-cell subset identification by Scissor

The workflow of Scissor is shown in Fig. 1. Denote *m* and *n* as the cell number and the bulk sample number, respectively. The two sources of data for Scissor inputs are a single-cell expression matrix with *m* cells and a bulk profiling data with sample phenotypes *Y*, which can be a binary group indicator vector or clinical survival data (Fig. 1a). Scissor first uses quantile normalization on single-cell and bulk expression data to remove the underlying batch effect. After this, a Pearson correlation matrix *S =* (*s*_*ij*_)_*n*×*m*_ is calculated for each pair of cell and bulk sample, where *s*_*ij*_ is the correlation of sample *i* and cell *j* across common genes in normalized single-cell and bulk expression data. Meanwhile, a corresponding cell-cell similarity network *G* is constructed based on single-cell data. In this study, we used the shared nearest neighbor graph embedded in Seurat^31^ to serve as *G*. Next, let *β* denotes a vector of coefficients and *l*(*β*) denotes an appropriately chosen log-likelihood function (The formula of *l*(*β*) depends on the type of phenotype (See more details in next section). Scissor integrates the correlation matrix *S*, sample phenotype *Y* and cell-cell similarity network *G* into a regression model by imposing penalties on *l*_1_-norm of coefficient and Laplacian regularization (Fig. 1b). The *l*_1_-norm penalty leads to a sparse cell selection result and the network-based penalty enforces that the tightly connected nodes (cells) in the network tend to have more similar coefficients^46^. Overall, Scissor is formulated as the following optimization model:

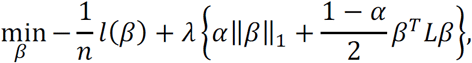

where *L* is the symmetric normalized Laplacian matrix, which is defined as

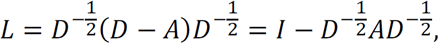

where *A =* (*a*_*ij*_)_*m*×*m*_ is the binary or weighted adjacency matrix of *G. a*_*ij*_ equals to 1 or a value ranging from 0 to 1, if cell *i* and *j* are connected in *G*, and *a*_*ij*_ *=* 0, otherwise. *D =* (*d*_*ij*_)_*m*×*m*_ is the degree matrix of *G*, where 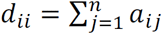, and *d*_*ij*_ *=* 0 for *i* ≠ *j*. The tuning parameter λ controls the overall strength of the penalty and *α* balances the amount of regularization for smoothness and sparsity.

The non-zero coefficients of *β* solved by the above optimization model are used to select cell subpopulations which are correlated with the phenotype of interest (Fig. 1c). According to the sign of *β*, we denote the selected cells by Scissor+ cells and Scissor-cells, respectively, and unselected cells are denoted as background cells. The association of Scissor+ and Scissor-cells with the phenotype depends on the phenotype input *Y*. For example, if we define the drug responder as 1 and non-responder as 0, the Scissor+ cells are positively correlated with the responder phenotype. Usually, Scissor+ cells and Scissor-cells can be largely unbalanced since they will capture the different natures of bulk phenotypes. Finally, these selected cells will further be passed to several downstream analyses, such as the differential gene expression analysis, functional enrichment analysis, motif analysis and so on, to reveal the biological mechanisms of the selected cell subpopulations (Fig. 1d).

### Detailed formula of log-likelihood function in Scissor

The selection of log-likelihood function *l*(*β*) in Scissor depends on the type of phenotype information *Y*. We have developed Scissor to address three types of *Y*: (1) linear regression for continuous clinical outcomes; (2) binomial logistic regression for the binary phenotype like the treatment response and nonresponse; (3) cox regression for survival information. Denote *S*_*i*_ *=* (*s*_*i*1_, *s*_*i*2_, …, *s*_*im*_)^*T*^ is the correlation coefficients for sample *i* across all *m* cells. If the continuous outcome *Y =* (*y*_1_, *y*_2_, …, *y*_*n*_) ^*T*^ is chosen to guide the cell selection, the following linear regression log-likelihood function is used:

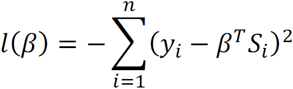

If *Y* is binary, e.g. *y*_*i*_ ∈ {0,1}, the binomial logistic regression is used:

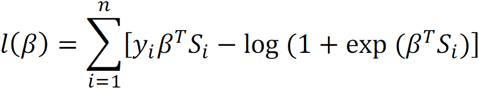

For time-to-event outcomes subject to independent censoring, the Cox model is considered. Let *T*_*i*_ be the non-negative event time and *C*_*i*_ be the censoring time. Denote 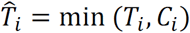 be the observed event time or censoring time and *δ*_*i*_ *= I*(*T*_*i*_ ≤ *C*_*i*_) be the event indicator, where *I*(.) is an indicator function. We use the following log-likelihood function:

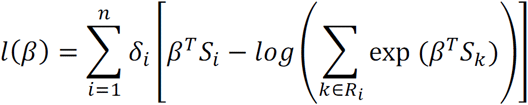

where 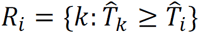 denotes the risk set at time 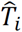.

### Parameter tunings and application details

In this study, the algorithm proposed by *Li et al*.^47^ is used to solve the above regularized regression model. In our model, there are two model parameters need to be determined. First, parameter λ controls the overall strength of the penalty. In Scissor, we set 100 possible values for λ and a 10-fold cross-validation is applied to select the optimal λ with the minimum cross-validation averaged error. Second, parameter *α* ∈ [0, 1] balances the effect of *l*_1_-norm and network-based penalties. Larger *α* inclines to lay more emphasis on *l*_1_-norm to encourage sparsity. Thus, Scissor will select fewer cells. In real applications, it is difficult to fix a default value of *α* for all kinds of datasets, since different datasets could have different sensitivities to the changes of *α*. Therefore, we made the following constraint on the total number of Scissor selected cells to determine *α*: the number of Scissor selected cells should not exceed the number of 10% of total cells in the single-cell data under study. In each experiment, a search on the default value list of *α*, {0.01, 0.05, 0.1, 0.2, 0.3, …, 0.9}, is performed from the smallest to the largest until a value of *α* meeting the above criteria.

There are in total three applications of Scissor in this study. In the tumor vs normal experiment and melanoma immune cell selection experiment, the logistic regressions with 0-1 binary response variables were used. The response variable equals to 1 for tumor/non-responder samples and 0 for normal/responder samples. In cancer cell selection experiment, the overall survival data was used and a Cox regression model was fitted in Scissor. For single-cell dataset with high dropout rate, it is necessary to perform imputation for the single-cell expression data. For our study, we used the imputed melanoma single-cell expression matrix as Scissor input, whereas kept the original expression of lung cancer single-cell data.

### Preprocessing single-cell RNA-seq data

In this study, we used the Seurat R package (version 3.1.2)^31^ to preprocess the original scRNA-seq data. After this, the preprocessed scRNA-seq expression matrix was fed into Scissor. As a quality control step, we first filtered out genes that are not expressed in at least 400 cells. If the input expression matrix is read counts data, the filtered expression matrix was normalized using the NormalizeData function with the ‘LogNormalize’ normalization method and ‘scale.factor’ equals to 10,000. Furthermore, we focused on the genes that exhibited high cell-to-cell variation and identified variable genes using the FindVariableFeatures function in Seurat with the default ‘vst’ method. Besides, principal components analysis was performed on the scaled data cut to variable genes and the first 10 principal components were selected for downstream clustering analysis. Cells were embedded in a shared nearest neighbor graph, which is the cell-cell similarity network used as the network-constrained regularization in Scissor. Cells were then partitioned into clusters using the FindClusters function with the resolution parameter set to 0.6 and the other parameters left as default. In order to visualize cells in low dimensions, uniform manifold approximation and projection (UMAP)^48^ was generated using the RunUMAP function.

### Differential gene expression analysis

For all single-cell differential gene expression tests, we used the default Wilcoxon rank-sum test implemented in Seurat. The differentially expressed genes for Scissor+ cells compared with all other cells (including Scissor-cells and background cells) were identified using the FindMarkers function. In this study, we required the following criteria to obtain the differential expression genes in each application: First, the gene expression between two groups is statistically significant with false discovery rate (FDR) < 0.05. Second, the absolute value of expression fold change between two groups is greater than 1.25. Last, the gene with expression value greater than 0 in at least 10% cells in either of the two groups was kept for further analysis. The VlnPlot function in Seurat was used to exhibit the relative expression levels of the selected differential expression genes.

Several signatures were generated based on the differential expression genes in each experiment. In lung cancer cells, the 23 overexpressed differential expression genes were defined as lung cancer signature and further proved its clinical relevance on six independent datasets. As for melanoma T cells, all 137 differential expression genes (17 up-regulated genes and 120 down-regulated genes) were claimed as our immunotherapy responsive signature.

In order to obtain a pre-ranked gene list (introduced below) that contains all candidate genes, we did not set thresholds for fold change (logfc.threshold = log(1)) and the minimum fraction of expression cells (min.pct = 0). The other parameters of FindMarkers were set to default.

### Pathway enrichment analysis

The pre-ranked version of camera (function name: cameraPR) compiled in the limma R package (version 3.38.3)^49^ was used to evaluate the enriched molecular pathways in Scissor+ cells. cameraPR needs a pre-ranked gene list according to a user-defined statistic score. In this study, we used the differential expression genes output of FindMarkers (ave_logFC and FDR) to obtain the pre-ranked gene list. In detail, the statistic ranking score for each gene was calculated using the following formula:

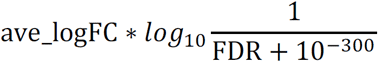

where ave_logFC stands for the log-transformed fold change between two groups and FDR is the adjusted Wilcoxon rank-sum test p-value. The gene sets were downloaded from the Molecular Signatures Database (MSigDB)^50^ (version 7.0). The Mognol_LCMV_Tumor_Exhaustion gene set contains the genes that are up-regulated in exhausted T cells from both OT-I mouse model and lymphocytic choriomeningitis virus (LCMV) infected mice, which was fetched from the Mognol’s study^51^.

### Survival analysis

For survival analysis, the gene set variation analysis (GSVA) algorithm with the default settings, as implemented in the GSVA R package (version 1.30.0)^24^ was applied to calculate the signature score for our lung cancer signature in each sample. Next, the samples were divided into two groups based on the quantile values of signature scores. Survival curves of these two groups of patients were estimated by the Kaplan-Meier method with statistical significance calculated using log-rank test. Furthermore, three other clinical features (pathological stage, sex and age at diagnosis) along with the lung cancer signature were evaluated by a univariate Cox proportional hazard (Cox PH) model to examine the association between survival time and these factors. The factors that were found to be prognostic significant (*p* < 0.05) in the univariate Cox PH model were included in the subsequent multivariable Cox PH model. The Kaplan-Meier survival analysis, log-rank test and Cox PH models were performed in the survival R package (version 3.1-8). When comparing the nested models (Cox PH model with tumor stage alone and the model with both tumor stage and lung cancer signature), the likelihood ratio test was used and achieved by the anova function in R.

### Motif analysis

The motif analysis for the signature genes was performed by oPOSSUM 3.0 software^52^ with the default parameters, which can detect the over-represented conserved transcription factor binding sites based on a set of gene names. The final motif list was ranked by the Fisher score.

### Gene set enrichment score for immunotherapy responsive signature

A pseudo-regulon was built based on our immunotherapy responsive signature: the 17 up-regulated genes and 120 down-regulated genes were served as positive and negative target genes of a pseudo-regulon, respectively. Then, the viper function compiled in the viper R package (version 1.16.0)^53^ was employed to infer the enrichment score of our signature.

### Gene set enrichment plot

The gene set enrichment plots of the 17 up-regulated genes and 120 down-regulated genes from the immunotherapy responsive signature were generated by clusterProfiler^54^ in Fig. 5h and Supplementary Figure 3.

### Pseudotime analysis

Slingshot^55^ with the default parameters was used to perform the pseudotime analysis for melanoma T cell dataset. The expressions of 137 genes in immunotherapy responsive signature across cells were used as the input of Slingshot.

### Lung cancer single-cell RNA-seq data

The lung cancer scRNA-seq data from *Lambrechts et al*. (ArrayExpress accessions E-MTAB-6149 and E-MTAB-6653)^11^ was used in this study. The expression data and the cell types were extracted from .loom source files using the loomR R package (version 0.2.0). The original scRNA-seq data contains 52,698 cells from 2 lung squamous carcinoma samples (4,314 cells), 2 LUAD samples (29,888 cells) and 1 non-small-cell lung cancer sample (18,496 cells). In order to make the cancer type of scRNA-seq data consistent with the corresponding bulk data and keep the majority of the total cells as much as possible, only cells from LUAD samples were used as scRNA-seq input of Scissor. In detail, the 29,888 cells containing malignant and non-malignant cells from LUAD patients were used in the identification of cell subpopulations related to tumor and normal phenotypes (Fig. 2a). Among the above 29,888 cells, 4,102 cells were marked as cancer cells in the original paper^11^ (Fig. 3a) and were analyzed by Scissor to identify a cancer cell subpopulation in LUAD associated with the TCGA-LUAD patient survival outcomes.

### Lung cancer bulk RNA-seq data

The lung cancer bulk RNA-seq Fragments Per Kilobase Million (FPKM) gene expression data and the clinical survival information used in this study were downloaded from TCGA-LUAD using the TCGAbiolinks R package (version 2.15.3)^56^. The FPKM was further converted into Transcripts Per Kilobase Million (TPM) quantification. The 577 samples with the tumor and normal phenotypes were used in the identification of cell subpopulations related to tumor and normal phenotypes. And the 471 TCGA-LUAD tumor samples with valid overall survival information were used to guide the cell subpopulation identification in the lung cancer cells.

### Lung cancer datasets for examining the clinical relevance of lung cancer signature

Six lung cancer datasets (GEO accession number: GSE13213, GSE11969, GSE31210, GSE8894, GSE3141 and GSE4573) with survival information were downloaded from the PRECOG website (https://precog.stanford.edu/index.php). These datasets were selected since their sample numbers are greater than 100. When mapping the probe ids to gene symbols, the median expression value of probe ids mapped to the same gene symbol was used.

### Melanoma single-cell RNA-seq data

The Tirosh’s melanoma scRNA-seq data^28^ was analyzed by Scissor to identify T cell subpopulations associated with immunotherapy. The preprocessed expression matrix was directly downloaded from GEO (accession number: GSE72056). Because of the established role of T cells in immunotherapy, we focused our analysis on 2,068 T cells labeled in the original paper. In our initial data inspection, three clusters were identified in this T cell dataset and the smallest cluster with 174 cells was characterized by the high expression of cell cycle-related genes. Another study also mentioned that these cells were contaminated with melanoma markers^34^. Therefore, we removed these cells and focused our analysis on the remaining 1,894 T cells. Besides, the T cell expression matrix was imputed using scRecover^57^ to address the dropout issue.

### Melanoma bulk RNA-seq data

70 melanoma patients with known immunotherapy response outcomes were collected from two studies^29, 30^. The Hugo’s cohort^29^ that contains 28 samples with 13 non-responders and 15 responders was downloaded from GSE78220. As for Allen’s cohort^30^, there are 42 samples with 28 non-responders and 14 responders, which were downloaded from the database of Genotypes and Phenotypes (dbGAP) under the accession number phs000452.v2.p1. The RNA-Seq by Expectation-Maximization (RSEM)^58^ was used to quantify the gene expression into TPM.

### Melanoma immunotherapy responsive signature validation in Sade-Feldman cohort

In Fig. 5h, the Sade-Feldman’s cohort^34^ was used as an independent data to test whether our immunotherapy responsive signature is informative in the prediction of CPI treatment success. The gene expressions of Sade-Feldman’s data were downloaded from GSE120575. All the gene expressions of cells for each patient were aggregated into one profile. Following the original paper^34^, the samples with *de novo* resistance to checkpoint therapy due to the complete loss of *B2M* or *HLA-A,B,C* were excluded in our study.

### Mouse liver tumor dataset for characterizing different T cell states

In Fig. 6a, five types of T cells from a mouse liver tumor model^38^ was used to explore the T cell states by calculating the enrichment score of our immunotherapy responsive signature. The raw fastq files were downloaded from GSE126973, which were analyzed by RSEM to quantify the gene expression in TPM.

### Memory-precursor-like and effector-like T cell data in Fig. 6c

The dataset that contains memory-precursor-like and effector-like T cell^40^ was used to characterize the Scissor selected T cells. The microarray gene expression data was downloaded from GSE8678, which profiled the Interleukin-7 Receptor (IL-7R) marked memory precursor effector CD8 T Cells and IL-7R low short-lived effector CD8 T cells sorted on day 6/7 after lymphocytic choriomeningitis virus (LCMV) infection in mice.

### Software availability

The open source Scissor program is freely available at GitHub https://github.com/sunduanchen/Scissor.

## Acknowledgements

This work was supported by the following funding: NIH 5K01LM012877 (to Z.X.); NIH 1R21HL145426 (to Z.X.); NIH 1R01CA207377 (to D.Z.Q); NIH NIGMS MIRA R35GM124704 (to A.A.); The Medical Research Foundation of Oregon (to Z.X.). We would like to thank Weston Anderson and Ann Hill for editing the manuscript. The resources of the Exacloud high performance computing environment developed jointly by OHSU and Intel and the technical support of the OHSU Advanced Computing Center are gratefully acknowledged.

## Author Contributions

D.S. and Z.X. conceived the idea, implemented the algorithm and performed the analyses. D.S., G.X., P.T.S. and Z.X. interpreted the results. A.M., D.Q., P.S. and A.A. provided scientific insights on the applications. Z.X. supervised the study. D.S. and Z.X. wrote the manuscript with feedback from all other authors. All of the authors read and approved the final manuscript.

## Competing interests

A.E. Moran discloses receipt of a sponsored research agreement from AstraZeneca. All other authors declare no competing interests.

## Supplementary figure legends

**Supplementary Figure 1.**
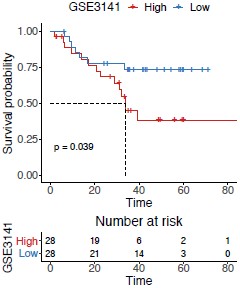
Lung cancer signature relates to prognostic information on the 5th validation datasets. Samples are divided into two groups based on the quantile values of lung cancer signature and Kaplan-Meier survival curve is drawn for each group. A corresponding table is also attached to display the number of alive samples at given time points and the statistical p value is calculated by log rank sum test.

**Supplementary Figure 2.**
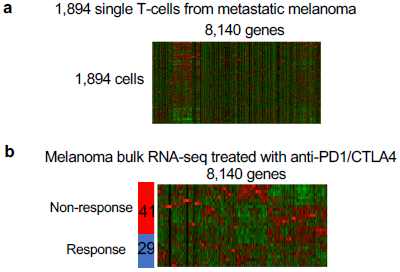
Inputs for the Scissor analysis of the scRNA-seq T-cells from the metastatic melanoma data. (**a**) The cartoon heatmap of 1,894 T-cell gene expressions from the scRNA-seq data of metastatic melanoma samples. (**b**) The gene expressions of 70 metastatic melanoma patients were obtained from the bulk RNA-seq. Those patients were treated with anti-PD1 or anti-CTLA4 from two studies with 41 non-responders and 29 responders.

**Supplementary Figure 3.**
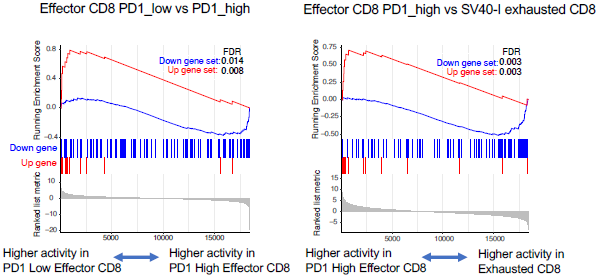
Gene set enrichment plot of the up and down immune signature genes in different CD8 T-cells. PD1 low Effector CD8: PD1low/LAG3low/CD39low tumor-specific CD8 T cells isolated Day 8-9 from malignant liver lesions. PD1 high Effector CD8: PD1high/LAG3high/CD39high TCR SV40-I CD8 T cells isolated Day 8-9 from malignant liver lesions. Exhausted CD8: TCR SV40-I CD8 T cells isolated Day 20-21 from malignant liver lesions. (**a**) The enrichment plot of our 137 immunotherapy responsive signature genes in the comparison between PD1 low effector CD8 and PD1 high effector CD8. (**b**) The enrichment plot of our 137 immunotherapy responsive signature genes in the comparison between PD1 high effector CD8 and Exhausted CD8 T cell.

## Supplementary table legends

**Supplementary Table 1**. The information of 23 up-regulated differential expression genes (lung cancer signature genes) in Scissor+ cells over all other cells (consisting of Scissor-cells and background cells). The Scissor+ cells were identified from lung cancer single-cell RNA-seq data, guided by the TCGA-LUAD bulk samples. The p-values were calculated by Wilcoxon rank-sum test.

**Supplementary Table 2**. The information of 205 down-regulated differential expression genes in Scissor+ cells over all other cells. The Scissor+ cells were identified from lung cancer single-cell RNA-seq data, guided by the TCGA-LUAD bulk samples. The p-values were calculated by Wilcoxon rank-sum test.

**Supplementary Table 3**. The full enriched motifs of 23 lung cancer signature genes. The motif analysis was performed by the oPOSSUM 3.0 software with default parameters. This motif list was ranked by the Fisher score.

**Supplementary Table 4**. The sample numbers and log-rank p-values of six lung cancer signature validation datasets. These datasets were downloaded from PRECOG and were selected since their sample numbers are larger than 100.

**Supplementary Table 5**. The information of 137 differential expression genes (immunotherapy responsive signature genes) in Scissor+ cells over all other cells. The Scissor+ cells were identified from melanoma T-cell single-cell RNA-seq data, guided by 70 melanoma bulk patient samples. The p-values were calculated by Wilcoxon rank-sum test.

